# A ComRS competence pathway in the oral pathogen *Streptococcus sobrinus*

**DOI:** 10.1101/2020.03.15.992891

**Authors:** Walden Li, Ryan M. Wyllie, Paul A. Jensen

## Abstract

*Streptococcus sobrinus* is one of two species of bacteria that cause dental caries (tooth decay) in humans. Our knowledge of *S. sobrinus* is limited despite the organism’s important role in oral health. It is widely believed that *S. sobrinus* lacks the natural competence pathways that are used by other streptococci to regulate growth, virulence, and quorum sensing. The lack of natural competence has also prevented genetic manipulation of *S. sobrinus*, limiting our knowledge of its pathogenicity.

We discovered a functional ComRS competence system in *S. sobrinus*. The ComRS pathway in *S. sobrinus* has a unique structure, including two copies of the transcriptional regulator ComR and a peptide pheromone (XIP) that lacks aromatic amino acids. We show that synthetic XIP allows transformation of *S. sobrinus* with plasmid or linear DNA, and we leverage this newfound genetic tractability to confirm that only one of the ComR homologs is required for induced competence.

Although *S. sobrinus* is typically placed among the mutans group streptococci, the *S. sobrinus* ComRS system is structurally and functionally similar to the competence pathways in the salivarius group. Like *S. salivarius*, the ComRS gene cluster in *S. sobrinus* includes a peptide cleavage/export gene, and the ComRS system appears coupled to a bacteriocin response system. These findings raise questions about the true phylogenetic placement of *S. sobrinus*.

Finally, we identified two strains of *S. sobrinus* appear to be “cheaters” by either not responding to or not producing XIP. While the mechanisms of cheating could be independent, we show how a recombination event in the non-responsive strain would restore function of the ComRS pathway but delete the gene encoding XIP. Thus the *S. sobrinus* ComRS pathway provides a lens into the evolution of ecological cheaters.

## Introduction

Dental caries (tooth decay) results from acid fermentation by two species of streptococci: *Streptococcus mutans* and *Streptococcus sobrinus* [1]. *S. mutans* is found in mouths of most of the world’s population, while the rarer *S. sobrinus* colonizes the teeth of 9%-15% of humans [2]. While both species are cariogenic by themselves, co-infection by *S. mutans* and *S. sobrinus* leads to aggressive caries and poorer oral health outcomes, especially in children [2, 3].

*S. sobrinus* is one of the few species of oral streptococci that lack a natural competence pathway. Competence is central to pathogenicity in streptococci, and competence pathways are connected to metabolism [4], virulence [5, 6], quorum sensing [7], and antibiotic tolerance [8]. The natural competence of streptococci has been known for over 75 years [9]. In 1995 it was discovered that a peptide pheromone named CSP controlled competence by signaling through the ComDE pathway [10, 11]. A second competence pathway, ComRS, was later discovered in many streptococci [12, 13]. ComRS systems contain a transcriptional regulator (ComR) that is activated by XIP, a small peptide derived from a precursor peptide ComS. ComRS systems are classified based on the sequence of the XIP peptide [12]. Type I XIPs are found in the salivarius group streptococci and have a characteristic P(F/Y)F motif. Type II XIPs contain a WW motif and are widespread in the bovis, pyogenes, and mutans groups, including the cariogenic species *S. mutans* [13], *S. ratti* [14], and *S. macacae* [15]. Some strains of *S. suis* contain Type II ComRS systems, but the majority have Type III XIPs with a WG(T/K)W motif. All known XIP sequences, regardless of type, contain two aromatic amino acids [15].

The classic XIP pathway functions as an autocrine loop (Figure 1A) [16]. The precursor peptide ComS is exported and cleaved to form active, extracellular XIP. The processed XIP is imported by an unknown mechanism and binds to the transcriptional regulator ComR. In *S. thermophilus* the XIP binding facilitates dimerization of ComR, and the ComR dimers bind a DNA motif to initiate transcription of competence genes [15].

Both the ComRS and ComDE systems activate the sigma factor ComX and other subsequent competence genes. *S. mutans* has functional ComDE and ComRS pathways, but genomic studies predict that both pathways are missing or incomplete in *S. sobrinus* [14, 17]. It is unclear how *S. sobrinus* can form communities and infect humans without a functional competence pathway.

**Figure 1.**
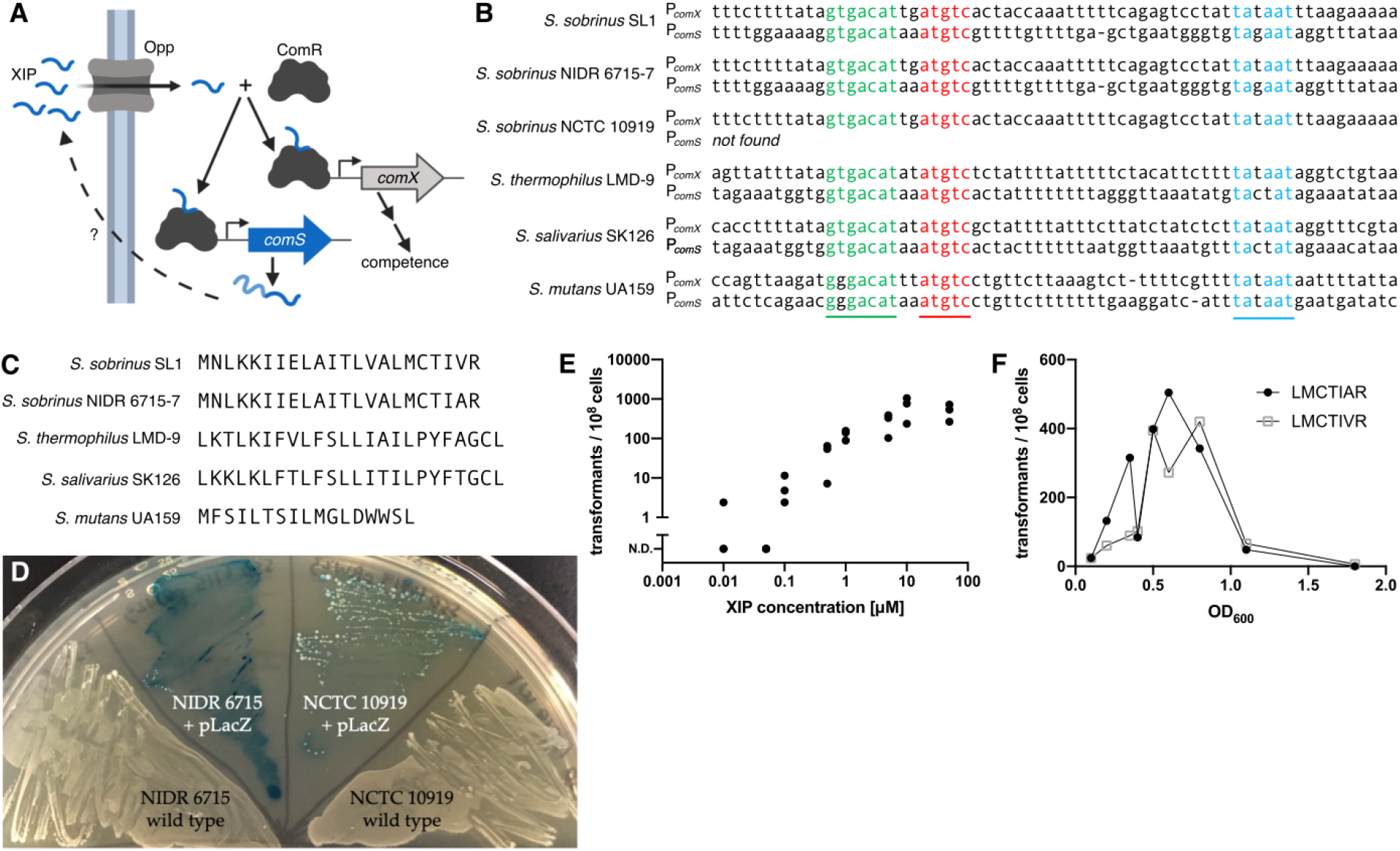
The peptide XIP induces competence in *S. sobrinus*. **A.** The ComRS competence pathway in *S. mutans* forms an autocrine signaling loop. An unknown protein cleaves the leader peptide from ComS and exports the XIP precursor. Activated XIP is imported where it facilitates dimerization of the transcriptional regulator ComR. The ComR/XIP complex binds a DNA motif to promote transcription of competence genes, including *comS*. **B.** The ComR/XIP binding motif appears upstream of the sigma factor *comX*. Using this sequence we identified a *comS* gene in three strains of *S. sobrinus*. The *comS* gene appears downstream of a homolog of the regulator *comR*. C. The *S. sobrinus* ComS peptides differ from sequences in *S. mutans S. salivarius*, and *S. thermophilus*. In particular, all XIP sequences in streptococci contain two aromatic amino acids; the XIP in *S. sobrinus* has none. **D.** *S. sobrinus* strains NIDR 6715-7 and NCTC 10919 can be transformed using exogenous XIP. Both strains were transformed with a plasmid expressing LacZ. When plated with X-gal, the plasmid-carrying strains produce a blue color, but the wild-type strains do not. E. The transformation efficiency of *S. sobrinus* strain NCTC 10919 increases with XIP concentration. Transformation assays used linear DNA with homology to regions flanking *comR*. No transformants without XIP. F. Transformation efficiency peaks in mid-exponential phase. Transformation assays were performed with strain NCTC 10919 and the pRW17 plasmid using the predicted XIP for strains SL1 (LMCTIVR) and NIDR 6715-7 (LMCTIAR). No XIP precursor gene (*comS*) is found in the NCTC 10919 genome.

The lack of natural competence also has practical effects on *S. sobrinus* research. *S. sobrinus* is widely believed to be genetically intractable. The genome of the well-studied *S. mutans* can be modified by hijacking the organism’s natural competence pathways [18–20]. Adding synthetic analogues of CSP or XIP causes *S. mutans* to readily uptake exogenous DNA. By contrast, there are no reports of induced competence in *S. sobrinus*. CSP and XIP from *S. mutans* do not induce competence in *S. sobrinus*, and other methods (i.e. horse serum or acid stress) fail to stimulate *S. sobrinus* like other oral streptococci. The paucity of genetic tools for *S. sobrinus* creates gaps in our mechanistic understanding of how this pathogen affects oral health.

Here we report that *S. sobrinus* does, in fact, contain a functional ComRS competence pathway. Although *S. sobrinus* is commonly classified among the mutans streptococci, the *S. sobrinus* ComRS system is similar to pathways in the salivarius group. Most strains of *S. sobrinus* contain two homologs of ComR, and we show that only one is essential for transformation with foreign DNA. Unlike all known XIP sequences, the XIPs from *S. sobrinus* contain no aromatic amino acids. Finally, we report two strains of *S. sobrinus* that appear to be “cheating” by not sensing or not producing XIP. One strain contains a truncated ComR and does not respond to exogenous XIP. A second strain can be transformed with XIP but has lost the genes encoding ComS and its exporter. Overall, the discovery of a novel competence pathway suggests that *S. sobrinus* can communicate extracellularly within the complex oral microbiome. The ComRS pathway also provides a robust method for genetic manipulation of *S. sobrinus*.

## Results

Previous studies have searched for ComDE and ComRS systems in *S. sobrinus* [14, 17]. Although some homologs of genes in the *S. mutans* ComDE or ComRS pathways were found in *S. sobrinus*, the pathways were largely incomplete and thought to be nonfunctional. Searches for ComS (XIP) and ComC (CSP) have also failed, although these genes are difficult to find. The *comS* gene is small (69 nucleotides) and is frequently missed by annotation pipelines. Genome mining in *S. sobrinus* has also been hampered by the lack of a complete genome sequence for any strain of *S. sobrinus*. The first complete genomes for *S. sobrinus* strains NIDR 6715-7, NCTC 10919, and the type strain SL1 were released in 2018 [21]. No search for *S. sobrinus* competence pathways has been published since the complete genomes became available.

We used a different approach to search for ComRS pathways in *S. sobrinus*. Rather than look for homologs of ComR, we searched for the promoter sequence recognized by the ComR/XIP complex. This ComR/XIP motif is located upstream of the *comX* gene in species with the ComRS pathway [13]. The ComR/XIP motif from *S. mutans* UA159 (GGGACATNNATGTC) was not present in *S. sobrinus*; however, the ComR/XIP motif from *S. thermophilus* LMD-9 and *S. salivarius* SK126 (GTGACATNNATGTC) was found upstream of a sigma factor homologous to *comX* in the genomes of *S. sobrinus* SL1, NIDR 6715-7,and NCTC 10919 (Figure 1B, Supplementary Table S1). In two of the *S. sobrinus* strains (SL1 and NIDR 6715-7), the ComR/XIP motif also appeared upstream of a short open reading frame. This ORF was downstream of a transcriptional regulator with homology to the *S. thermophilus comR* gene, suggesting that the ORF may be *comS* (Supplementary Table S1).

The 22 amino acid ComS peptide is similar in the NIDR 6715-7 and SL1 strains of *S. sobrinus* (Figure 1C). Both sequences differ from the ComS peptides in *S. thermophilus* and *S. salivarius* and lack a tryptophan-tryptophan sequence near the C-terminus that is characteristic of ComS in the mutans streptococci [13, 16]. We were surprised that the *S. sobrinus* ComS sequence contains no aromatic amino acids since all other streptococcal XIP sequences have two aromatic residues that interact with ComR during binding [15]. We predicted that the mature XIP in *S. sobrinus* would be the final seven amino acids of ComS based on the cleavage patterns in *S. thermophilus* and *S. salivarius*. We purchased synthetic peptides and tested their ability to induce competence in *S. sobrinus* by adding plasmid DNA and selecting for transformants. The XIP peptides from strains NIDR 6715-7 and SL1 induced competence in strain NIDR 6715-7; we were unable to transform strain SL1 with either peptide (Figure 1D). The genome of strain NCTC 10919 does not contain a *comS* gene, but the strain is transformable using XIP from either NIDR 6715-7 or SL1 (Figure 1D). In addition to plasmid DNA, we were able to transform *S. sobrinus* using linear DNA via homologous recombination. Strains NIDR 6715-7 and NCTC 10919 can also be transformed using a synthetic peptide with the last nine amino acids of NIDR 6715-7 ComS. We did not observe any transformants using a shorter peptide with only five amino acids.

The transformation efficiency of *S. sobrinus* increases with XIP concentration, peaking at 10^−5^ transformants per cell with 10 μM XIP for strain NIDR 6715-7 (Figure 1E). The transformation efficiency also depends on the growth phase of the cells. The efficiency is highest in mid-exponential phase, similar to other streptococci transformed with XIP [19, 20].

In *S. sobrinus* the *comS* gene is located upstream of a predicted amino acid cleavage and export protein (Figure 2A, Supplementary Table S2). This arrangement is similar to *comS* gene cluster in *S. thermophilus* and *S. salivarius*, where the export/cleavage protein removes the ComS leader sequence and exports the XIP precursor [22]. The *comS* gene cluster in *S. mutans* does not contain the export/cleavage protein (Figure 2A).

**Figure 2.**
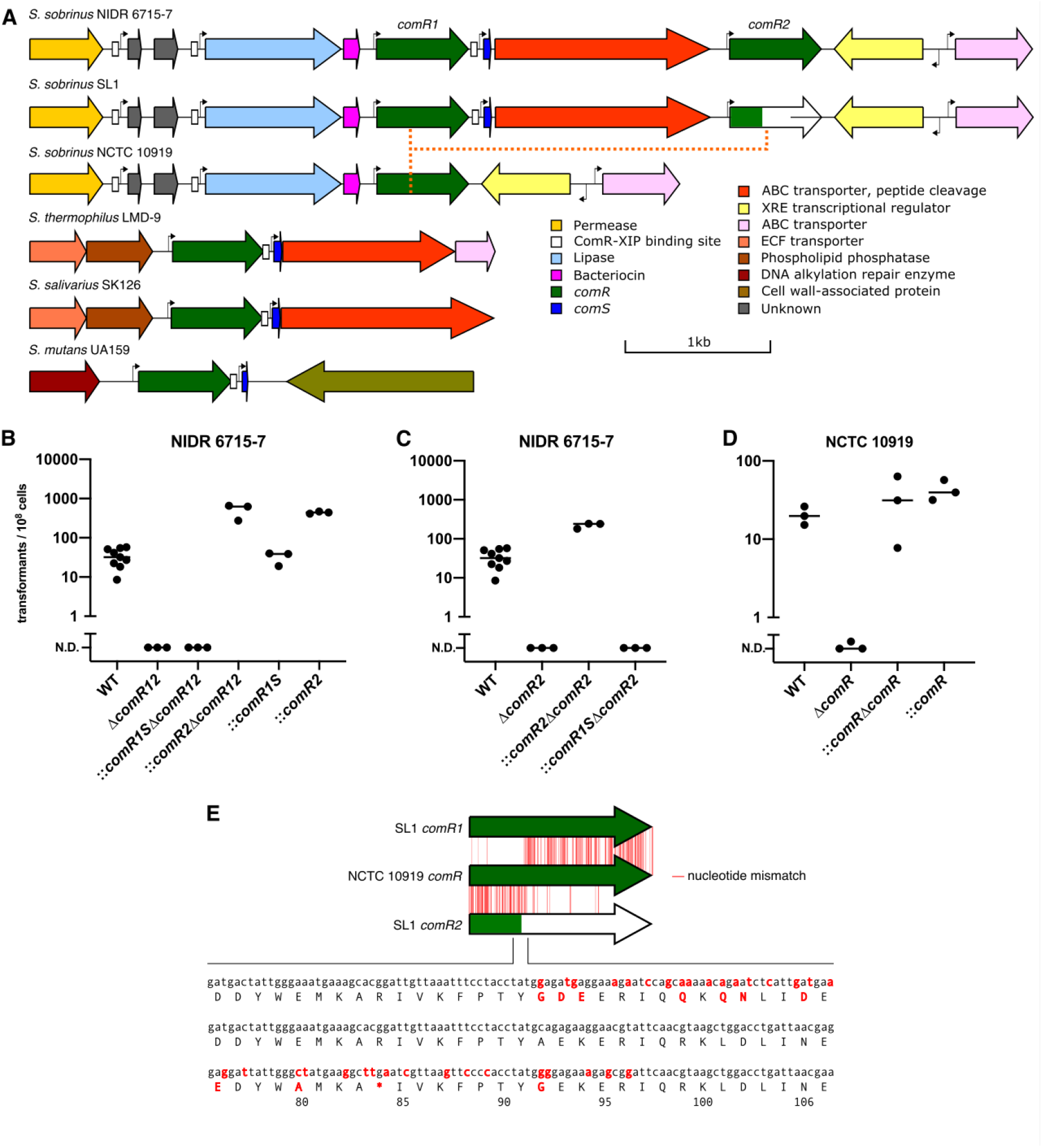
The *S. sobrinus* ComRS gene cluster is distinct from other streptococci. **A.** The ComRS gene cluster in strains NIDR 6715-7 and SL1 contain two homologs of *comR* that we call *comR1* and *comR2*. The type strain SL1 contains a truncated *comR2* gene (green/white) and cannot be transformed. Strain NCTC 10919 has a single homolog of *comR* and no *comS* gene or a ComS export/cleavage gene. **B.** Strain NIDR 6715-7 cannot be transformed if the region from *comR1* to *comR2* is deleted. Additional copies of *comR1* and *comS* on a plasmid to not rescue transformation, but strains complemented with extra *comR2* can be transformed. The horizontal bars represent the mean of the biological replicates (black dots). **C.** Loss of only the *comR2* gene prevents transformation of strain NIDR 6715-7. Transformability can be restored with additional copies of *comR2* but not *comR1*. **D.** The single *comR* homolog in strain NCTC 10919 is required for transformation. **E.** The *comR* gene in strain NCTC 10919 appears to be a fusion of *comR1* and *comR2*. A recombination event that produced *comR* would have removed the premature stop codon found in the *comR2* gene of strain SL1.

The structure of the *comS* gene cluster varies across the three *S. sobrinus* strains (Figure 2A, Supplementary Table S2). Strain NIDR 6715-7 contains two homologs of *comR* that we call *comR1* and *comR2* (Figure 2A). These genes flank the *comS* gene and the gene encoding the putative ComS exporter. The *S. sobrinus* type strain SL1 also contains *comR1* and *comR2*; however, *comR2* is truncated by a premature stop codon after 84 of 301 amino acids. Strain NCTC 10919 contains only a single *comR* gene and neither *comS* nor a gene encoding an export/cleavage protein.

We tested if *comR1*, *comR2*, or both genes are required for transformation. We constructed a Δ*comR12* strain from NIDR 6715-7 by deleting *comR1*, *comR2*, and the two genes in between (*comS* and the exporter) via homologous recombination. We tested if the deletion strain was transformable by attempting to delete a gene encoding a lactate oxidase (*lox*, DLJ51_01765 in NIDR 6715-7 and DK181_01730 in NCTC 10919) in the Δ*comR12* background. The Δ*comR12* strain could not be subsequently transformed using XIP and linear or plasmid DNA (Figure 2B). We were unable to perform a traditional complementation experiment since the Δ*comR12* strain cannot be transformed. Instead, we added plasmids carrying either *comR1* and the adjacent *comS* or *comR2* to the NIDR 6715-7 wild type strain and then reconstructed the Δ*comR12* mutation. The extra copies of *comR1* and *comS* do not complement the untransformable phenotype of the Δ*comR12* strain (Figure 2B). By contrast, strains with extra copies of *comR2* can be transformed with after adding Δ*comR12* mutation (Figure 2B). These data suggest that *comR2*, but not *comR1*, is required for transformation using exogenous XIP. We verified our results by deleting and complementing *comR2* alone (Figures 2C & 2D). In all cases, strains with at least one copy of *comR2* are transformable independent of the presence or absence of *comR1*.

Strains of NIDR 6715-7 carrying additional copies *comR1* or *comR2* on plasmids have different transformation efficiencies than the wild type strain (Figure 2B). We used a linear statistical model to quantify the overall effects of deleting or complementing *comR1* and *comR2*. Deleting *comR2* causes a significant reduction in transformation efficiency (p < 5 × 10^−11^). On average, strains of NIDR 6715-7 with extra copies of *comR2* in *trans* are transformable with 10-fold higher efficiency (p < 1.5 × 10^−9^). The effect of modifying *comR1* is less clear. The main effect of deleting *comR1* is a 5.2-fold increase in transformation efficiency (p < 6 × 10^−5^), but strains carrying extra copies of *comR1* also show increased transformation efficiency (2.1-fold, p < 0.014).

We performed similar deletion and complementation experiments in *S. sobrinus* NCTC 10919, the strain with one *comR* homolog and no *comS*. The Δ*comR* strains were largely untransformable (Figure 2E). We once observed a single colony on a plate but were unable to determine if this colony was a true transformant; we have never seen a transformant since. A strain of NCTC 10919 with additional copies of *comR* in *trans* retains competence after the original copy of *comR* is deleted from the chromosome (Figure 2E). We conclude that the single homolog of *comR* in *S. sobrinus* NCTC 10919 is necessary for XIP-induced competence.

The importance of *comR2* for competence may explain the intractability of strain SL1 and the single homolog of *comR* in strain NCTC 10919. Based on the structure of *comR* in *S. thermophilus* [15], we predict that the truncated *comR2* in strain SL1 is not functional, and thus the SL1 strain should not be competent. It appears that the single copy of *comR* is the result of a recombination event between *comR1* and *comR2* since the 5’ end of the gene is more similar to *comR1* and the 3’ end is more similar to *comR2* (Figure 2F). The putative recombination site would have removed the premature stop codon at amino acid 84 in the *comR2* of strain SL1. The *comR1*/*comR2* recombination in NCTC 10919 could be independent of the *comR2* truncation in SL1. It is also possible that the recombination restored function of the broken ComRS pathway in a common ancestor of the extant strains.

The ComR boxes recognized by the ComR/XIP complex are canonically associated with the promoters of *comX* and *comS*. In *S. salivarius* and *S. thermophilus*, ComR boxes are also found upstream of multiple bacteriocin gene clusters. The bacteriocins in *S. salivarius* are broad-spectrum antibacterials and are coupled to XIP-induced transcriptional activation [23].

We identified two additional ComR boxes in the SL1, NIDR 6715-7, and NCTC 10919 strains of *S. sobrinus*. Both boxes are near the ComRS gene cluster (Figure 2A) and are located 25 bp upstream of a sequence matching the consensus −10 promoter element (TATAAT). The first ComR box is upstream of an unannotated 42 amino acid peptide containing a double-glycine motif. The motif in *S. sobrinus* (LTxxDLxxVxGG) is similar to a known motif (LSxxELxxIxGG) [24] that is associated with peptide secretion, quorum-sensing, and bacteriocin production systems in streptococci [25]. The second ComR box in *S. sobrinus* is upstream of a predicted lipase (Figure 2A). The lipase gene is followed by a 25 bp intergenic region and a punitive bacteriocin. This bacteriocin also contains a double-glycine motif (LNxxDLxxIxGG).

The ComRS pathway is widespread among strains of *S. sobrinus*. A BLAST search revealed that all of the 54 publicly available genomes for *S. sobrinus* contain a homolog of the *comR* gene. In 83% of these genomes the ComR amino acid sequence has greater than 97% amino acid identity with ComR2 in *S. sobrinus* NIDR 6715-7. The conservation of ComR suggests that XIP may be useful for transforming a wide range of *S. sobrinus* strains.

## Discussion

A functional competence pathway changes our ecological view of *S. sobrinus*. The ComRS is used by other streptococci for inter- and intracellular communication, and now know that *S. sobrinus* could participate in these community-level conversations. The structure of the *S. sobrinus* ComRS gene cluster is unique among streptococci, and the lack of aromatic amino acids in the XIP sequence is novel. The ComRS systems most similar to *S. sobrinus* are found in the salivarius group (*S. salivarius* and *S. thermophilus*). *S. sobrinus* was originally a subspecies of *S. mutans* and is frequently grouped with the mutans streptococci [14]. Recent phylogenetic trees place *S. sobrinus* closer to *S. salivarius* and *S. thermophilus* or in a separate sobrinus clade [26]. Our functional data on ComRS would agree with *S. sobrinus*’s placement in or near the salivarius group. However, neither *S. salivarius* or *S. thermophilus* are cariogenic, raising questions about the association between *S. sobrinus* infection and aggressive disease.

Activating the ComRS pathway using synthetic XIP provides an efficient method for transforming *S. sobrinus*. We believe induced competence will open new avenues of *S. sobrinus* research by allowing forward genetic studies. We are not the first group to report genetic manipulation of *S. sobrinus*. One publication claimed that *S. sobrinus* can be transformed via electroporation [27], but we and other groups failed to reproduce this result [14]. In 1995 LeBlanc et al. reported the knockout of the *scrA* gene in *S. sobrinus* strain NIDR 6715-10 using a conjugation-based system [28]. We reconstructed the conjugation system and were able to transfer plasmids into multiple species in the mutans group, but not *S. sobrinus*. Interestingly, we could not find a homolog of the *scrA* gene deleted by LeBlanc in any *S. sobrinus* genome. However, the reported *scrA* sequence [29] has >99% identity to a gene in *S. ferus*. We were unable to find the original NIDR 6715-10 strain used by LeBlanc, so we cannot confirm if the strain was truly *S. sobrinus*. Regardless, we now have an easier method of transforming *S. sobrinus* using via XIP-induced competence.

The ComRS gene clusters in our *S. sobrinus* strains may indicate two methods of “cheating” [12]. The truncation of *comR2* in strain SL1 makes it unresponsive to XIP *in vitro*. It is possible that strain SL1 responds to XIP *in vivo*, especially since we do not understand the function of *comR1*. The ComR boxes upstream of bacteriocins in *S. sobrinus* suggest that XIP may induce an antibiotic defense response. A non-functional ComR2 could allow SL1 to opt-out of its share of the community defense. There are multiple explanations for the loss of *comS* by recombination in strain NCTC 10919. The strain still responds to exogenous XIP, so cells could save the cost of producing XIP while sensing XIP from other strains of *S. sobrinus*. Alternatively, NCTC 10919 could descend from a strain of *S. sobrinus* that shared the truncated *comR2* gene. The recombination event would have removed the premature stop codon and restored the ability to sense XIP. The loss of *comS* and its exporter would be a tradeoff for restoring the function of ComR.

Many streptococci produce bacteriocins, but a direct link between XIP and bacteriocins has only been observed in the salivarius group [25]. In *S. mutans*, bacteriocin production is activated indirectly through the ComDE system via ComX [30]. The ComR boxes upstream of bacteriocins in *S. sobrinus* suggest that XIP may induce an antimicrobial defense similar to *S. thermophilus* and *S. salivarius*. This is further evidence that *S. sobrinus*, or at least its competence pathway, is functionally closer to the salivarius group streptococci.

## Methods

### Strains, reagents, and growth conditions

Supplementary Table S3 lists the strains and plasmids used in this study. Liquid cultures for transformation assays were grown anaerobically (5% H_2_, 10% CO_2_, and 85% N%2) at 37°C in chemically defined medium (CDM) [31] containing 1% glucose. Solid agar plates were made with Todd-Hewitt broth plus 0.5% yeast extract, 1.5% agar, and antibiotic selection when needed. Final antibiotic concentrations were 1 mg/ml kanamycin, 400 μg/ml spectinomycin, and 4 μg/ml chloramphenicol. Plates were incubated aerobically under 5% CO_2_ at 37°C. All chemicals were purchased from Sigma Aldrich (St. Louis, USA) unless otherwise stated. Enzymes were purchased from New England Biolabs (Ipswich, MA, USA). Oligonucleotides were synthesized by Integrated DNA Technologies (Coralville, IA, USA). Synthetic peptides were purchased from GenScript, Inc. (Piscataway, NJ, USA) at >90% purity.

### Confirming transformants

Initial transformation assays used a custom *E. coli*/streptococcal shuttle vector carrying *lacZ* under the P23 broad-range Gram positive promoter (Supplementary Methods). Strains carrying the pLacZ plasmid appear blue when plated on 0.008% X-gal (5-bromo-4-chloro-3-indolyl β-D-galactopyranoside) (Zymo Research, CA, USA). X-gal concentrations above 0.008% appear to inhibit growth of *S. sobrinus* expressing LacZ. Knockout strains were confirmed by PCR using primers within the antibiotic marker and outside the homology arms of the linear DNA. The amplicons were Sanger sequenced to confirm the location of the knockout cassette.

### DNA manipulation and strain construction

Plasmids and linear DNA fragments were constructed by Golden Gate assembly [32, 33] using PCR amplicons and the primers in Supplementary Table S4. For gene deletions, approximately 1 kb of genomic DNA was amplified upstream and downstream of the gene of interest. These homology arms were digested with BsaI and joined to an antibiotic resistance cassette with T4 ligase. Antibiotic selection markers were *aph3* (kanamycin, [34]); *cat* (chloramphenicol, [35]), and *aad9* (spectinomycin, [36]). The genomic coordinates of each deletion site are reported in Supplementary Table S3. For plasmids, a new Golden Gate compatible backbone (pRW17) was constructed from the *E. coli*/streptococcal shuttle vector pDL278 [36] (Supplementary Methods). Final plasmids were assembled by one-pot digestion/ligation with BsaI and T4 ligase.

### Transformation assays

An overnight culture of *S. sobrinus* in CDM with antibiotic selection was diluted 150X to fresh CDM without antibiotics. The culture was grown to mid-exponential phase (OD600 between 0.55-0.75). The seven amino acid XIP peptide (disolved in DMSO) was added to a final concentration of 10 μM. The culture was mixed well by flicking the tube and 200 μl was transferred to a microcentrifuge tube where 400ng of DNA was added. After another 2h anaerobic incubation, 150 μl was plated on solid agar and incubated for 24 h aerobically. Transformation efficiency was calculated by comparing colony counts after 24 h on selective and nonselective plates. A detailed transformation protocol is available in the Supplementary Methods.

### Bioinformatic searches

Targeted queries for ComRS homologs were performed via BLAST [37] with blastp or tblastn through the NCBI web interface. Searches for homologs of ComR in all *S. sobrinus* strains was performed offline using custom R scripts and the blastp toolkit [38].

### Statistical analysis

Data were analyzed using Graphpad Prism or the R programming language. Inferences about the effects of *comR* deletion and complementation were based on t-tests of regression coefficients in a linear model. The model predicts log-transformation efficiency based on the main effects and interactions of *comR1* and *comR2* at three levels: knockout, wild type, and complemented. The true transformation efficiency is unknown for any strain that fails to produce colonies. For modeling we assume a single colony was observed as an upper bound on the transformation efficiency.

## Acknowledgements

We thank Bill Metcalf, Jim Slauch, and the members of the Jensen Lab for insightful discussions. We also thank Will Herbert and Emma Lee for assisting with the bioinformatic searches. This work was supported by the National Institutes of Health grant DE026817.

