## Supplementary Information for "A ComRS competence pathway in the oral pathogen *Streptococcus sobrinus*"

### Supplementary Table S1. Selected NCBI BLAST (tblastn) results of relevant ComR homologs

| Template | Species, Strain | Locus ID | Punitive Function(?) | Chromosome Indexing | Query Coverage | Coverage Indexing | E-Value | % Identity |
|---|---|---|---|---|---|---|---|---|
| SSO_SL1_ComR1 | SSO_SL1 | DK182_00280 | ComR2 | 43319-44221 | 99% | 43319-44218 | 7e-131 | 72% (217/300) |
| | SSO_6715-15 | DLJ52_00265 | ComR1 | 38804-39709 | 100% | 38804-39709 | 0.0 | 99 % (298/302) |
| | SSO_6715-15 | DLJ52_00275 | ComR2 | 42279-43178 | 99% | 42279-43178 | 4e-132 | 72% (217/300) |
| | SSO_10919 | DK181_00265 | ComR | 38716-39618 | 99% | 38716-39615 | 5e-146 | 79% (237/300) |
| | SMU_UA159 | SMU_381c | Unknown | 359781-360683 | 99% | 359784-360683 | 3e-127 | 75% (226/300) |
| | SMU_UA159 | SMU_61 | ComR | 61631-62545 | 60% | 61640-62173 | 7e-29 | 37% (67/179) |
| | STL_LMD-9 | STER_0316 | ComR | 272013-272912 | 99% | 272013-272909 | 6e-63 | 41% (122/299) |
| | SSL_NCTC8618 | NCTC8618_02749 | ComR | 319683-320582 | 99% | 319683-320579 | 2e-58 | 42% (125/299) |
| SSO_SL1_ComR2 | SSO_SL1 | DK182_00270 | ComR1 | 39843-40748 | 99% | 39843-40742 | 2e-120 | 72% (217/300) |
| | SSO_6715-15 | DLJ52_00275 | ComR2 | 42279-43178 | 99% | 42279-43181 | 2e-173 | 99% (299/301) |
| | SSO_6715-15 | DLJ52_00265 | ComR1 | 38804-39709 | 99% | 38804-39703 | 2e-119 | 71% (214/300) |
| | SSO_10919 | DK181_00265 | ComR | 38716-39618 | 100% | 38716-39618 | 6e-154 | 90% (270/301) |
| | SMU_UA159 | SMU_381c | Unknown | 359781-360683 | 100% | 359781-360683 | 4e-109 | 70 % (210/301) |
| | SMU_UA159 | SMU_61 | ComR | 61631-62545 | 60% | 61643-62176 | 3e-20 | 34% (61/178) |
| | STL_LMD-9 | STER_0316 | ComR | 272013-272912 | 99% | 272013-272909 | 2e-57 | 41% (123/299) |
| | SSL_NCTC8618 | NCTC8618_02749 | ComR | 319683-320582 | 99% | 319683-320579 | 1e-52 | 42% (125/299) |
| | SSL_NCTC8618 | NCTC8618_02909 | DNA-binding protein | 483624-484547 | 55% | 483981-484463 | 1e-9 | 30% (49/164) |
| SSO_6715-15_ComR1 | SSO_6715-15 | DLJ52_00275 | ComR2 | 42279-43178 | 100% | 42279-43178 | 2e-130 | 71% (214/300) |
| | SSO_SL1 | DK182_00270 | ComR1 | 39843-40748 | 100% | 39843-40748 | 0.0 | 99% (298/302) |
| | SSO_SL1 | DK182_00280 | ComR2 | 43319-44221 | 100% | 43319-44218 | 5e-129 | 71% (214/300) |
| | SSO_10919 | DK181_00265 | ComR | 38716-39618 | 100% | 38716-39615 | 2e-146 | 79% (237/300) |
| | SMU_UA159 | SMU_381c | Unknown | 359781-360683 | 99% | 359784-360683 | 1e-128 | 76% (227/300) |
| | SMU_UA159 | SMU_61 | ComR | 61631-62545 | 60% | 61640-62173 | 3e-29 | 38% (68/179) |
| | STL_LMD-9 | STER_0316 | ComR | 272013-272912 | 99% | 272013-272909 | 2e-63 | 41% (123/299) |
| | SSL_NCTC8618 | NCTC8618_02749 | ComR | 319683-320582 | 99% | 319683-320579 | 6e-59 | 42% (126/299) |
| SSO_6715-15_ComR2 | SSO_6715-15 | DLJ52_00265 | ComR1 | 38804-39709 | 100% | 38804-39703 | 1e-120 | 71% (214/300) |
| | SSO_SL1 | DK182_00280 | ComR2 | 43319-44221 | 100% | 43319-44221 | 2e-173 | 99% (300/301) |
| | SSO_SL1 | DK182_00270 | ComR1 | 39843-40748 | 100% | 39843-40741 | 9e-122 | 72% (217/300) |
| | SSO_10919 | DK181_00265 | ComR | 38716-39618 | 100% | 38716-39618 | 1e-155 | 90% (272/301) |
| | SMU_UA159 | SMU_381c | Unknown | 359781-360683 | 100% | 359781-360683 | 1e-109 | 69% (209/301) |
| | SMU_UA159 | SMU_61 | ComR | 61631-62545 | 60% | 61643-62176 | 5e-20 | 34% (60/178) |
| | STL_LMD-9 | STER_0316 | ComR | 272013-272912 | 99% | 272013-272909 | 5e-59 | 42% (125/299) |
| | SSL_NCTC8618 | NCTC8618_02749 | ComR | 319683-320582 | 99% | 319683-320579 | 2e-54 | 42% (127/299) |

| | | | | | | | | |
|---|---|---|---|---|---|---|---|---|
| | SSL_NCTC8618 | NCTC8618_02909 | DNA-binding protein | 483624-484547 | 95% | 483639-484532 | 3e-11 | 24% (77/317) |
| SSO_10919_ComR | SSO_6715-15 | DLJ52_00275 | ComR2 | 42279-43178 | 99% | 42279-43181 | 1e-163 | 90% (272/301) |
| | SSO_6715-15 | DLJ52_00265 | ComR1 | 38804-39709 | 100 | 38804-39703 | 5e-144 | 79% (237/300) |
| | SSO_SL1 | DK182_00280 | ComR2 | 43319-44221 | 100% | 43319-44221 | 4e-162 | 90% (270/301) |
| | SSO_SL1 | DK182_00270 | ComR1 | 39843-40748 | 99% | 39843-40742 | 1e-143 | 79% (237/300) |
| | SMU_UA159 | SMU_381c | Unknown | 359781-360683 | 100% | 359781-360683 | 2e-125 | 73% (220/301) |
| | SMU_UA159 | SMU_61 | ComR | 61631-62545 | 60% | 61640-62161 | 1e-24 | 34% (60/175) |
| | STL_LMD-9 | STER_0316 | ComR | 272013-272912 | 99% | 272013-272909 | 5e-61 | 42% (125/299) |
| | SSL_NCTC8618 | NCTC8618_02749 | ComR | 319683-320582 | 99% | 319683-320579 | 1e-56 | 43% (128/299) |

### Supplementary Table S2. Genomic architecture of ComRS locus in selected organisms

| Organism | Element Name | Size | Strand | Gene Name or Punitive Function |
|---|---|---|---|---|
| *Streptococcus sobrinus* SL1 | DK182_00250 | 720 | + | Permease |
| | comrxip1 | 20 | + | ComR-XIP binding site |
| | promoter1 | 10 | + | Promoter |
| | peptide | 130 | + | Unknown, unannotated |
| | DK182_00255 | 231 | + | Unknown |
| | comrxip2 | 20 | + | ComR-XIP binding site |
| | promoter2 | 10 | + | Promoter |
| | DK182_00260 | 1338 | + | Lipase |
| | DK182_00265 | 159 | + | Bacteriocin |
| | promoter3 | 10 | + | Promoter |
| | DK182_00270 | 906 | + | ComR |
| | comrxip3 | 20 | + | ComR-XIP binding site |
| | promoter4 | 10 | + | Promoter |
| | ComS | 70 | + | ComS |
| | DK182_00275 | 2118 | + | ABC transporter, peptide cleavage |
| | promoter5 | 10 | + | Promoter |
| | DK182_00280 | 903 | + | XRE transcription factor |
| | DK182_00285 | 873 | - | XRE transcription factor |
| | promoter6 | 10 | - | Promoter |
| | promoter7 | 10 | + | Promoter |
| | DK182_00290 | 759 | + | ABC transporter |
| *Streptococcus sobrinus* 6715-15 | DLJ52_00245 | 720 | + | Permease |
| | comrxip1 | 20 | + | ComR-XIP binding site |
| | promoter1 | 10 | + | Promoter |
| | peptide | 129 | + | Unknown, unannotated |
| | DLJ52_00250 | 231 | + | Unknown |
| | comrxip2 | 20 | + | ComR-XIP binding site |
| | promoter2 | 10 | + | Promoter |
| | DLJ52_00255 | 1338 | + | Lipase |
| | DLJ52_00260 | 159 | + | Bacteriocin |
| | promoter3 | 10 | + | Promoter |
| | DLJ52_00265 | 906 | + | ComR |
| | comrxip3 | 20 | + | ComR-XIP binding site |
| | promoter4 | 10 | + | Promoter |
| | ComS | 72 | + | ComS |
| | DLJ52_00270 | 2118 | + | ABC transporter, peptide cleavage |
| | promoter5 | 10 | + | Promoter |
| | DLJ52_00275 | 903 | + | XRE transcription factor |
| | DLJ52_00280 | 873 | - | XRE transcription factor |
| | promoter6 | 10 | - | Promoter |
| | promoter7 | 10 | + | Promoter |
| | DLJ52_00285 | 759 | + | ABC transporter |
| *Streptococcus sobrinus* 10919 | DK181_00245 | 720 | + | permease |
| | comrxip1 | 20 | + | ComR-XIP binding site |
| | promoter1 | 10 | + | Promoter |
| | peptide | 129 | + | Unknown, unannotated |
| | DK181_00250 | 231 | + | Unknown |
| | comrxip2 | 20 | + | ComR-XIP binding site |
| | promoter2 | 10 | + | Promoter |
| | DK181_00255 | 1338 | + | Lipase |
| | DK181_00260 | 159 | + | Bacteriocin |
| | promoter3 | 10 | + | Promoter |
| | DK181_00265 | 903 | + | ComR |
| | DK181_00270 | 873 | - | XRE transcription factor |
| | promoter4 | 10 | - | Promoter |
| | promoter5 | 10 | + | Promoter |
| | DK181_00275 | 759 | + | ABC transporter |
| *Streptococcus thermophilus* LMD-9 | STER_0314 | 558 | + | ECF transporter |
| | STER_0315 | 651 | + | Phospholipid phosphatase |
| | promoter1 | 10 | + | Promoter |
| | STER_0316 | 900 | + | ComR |
| | comrxip1 | 20 | + | ComR-XIP binding site |
| | promoter2 | 10 | + | Promoter |

| | | | | |
|---|---|---|---|---|
| | ComS | 78 | + | ComS |
| | STER_0317 | 1697 | + | ABC transporter, peptide cleavage |
| | STER_0318 | 390 | + | ABC transporter |
| *Streptococcus salivarius* SK126 | STRSA0001_1792 | 558 | + | ECF transporter |
| | STRSA0001_1791 | 651 | + | Phospholipid phosphatase |
| | promoter1 | 10 | + | Promoter |
| | STRSA0001_1790 | 900 | + | ComR |
| | comrxip1 | 20 | + | ComR-XIP binding site |
| | promoter2 | 10 | + | Promoter |
| | ComS | 78 | + | ComS |
| | STRSA0001_1789 | 2097 | + | ABC transporter, peptide cleavage |
| *Steptococcus mutans* UA159 | SMU_60 | 687 | + | DNA repair enzyme |
| | promoter1 | 10 | + | Promoter |
| | SMU_61 | 915 | + | ComR |
| | comrxip1 | 20 | + | ComR-XIP binding site |
| | promoter2 | 10 | + | Promoter |
| | ComS | 54 | + | ComS |
| | SMU_63c | 1842 | - | Cell-wall associated protein |

**Supplementary Table S3. List of Strains and Plasmids**

| Strain or plasmid | Characteristic(s) | Source |
|---|---|---|
| ***S. sobrinus*** | | |
| NCTC 10919 | Wild type | ATCC 33402 |
| NIDR 6715-7 | Wild type | ATCC 27351 |
| SL1 (DSM 20742) | Wild type | ATCC 33478 |
| NCTC 10919::*comR* | NCTC 10919 pComR | This study |
| NCTC 10919::*comR*Δ*comR* | NCTC 10919 Δ*comR::cat* pComR | This study |
| NCTC 10919Δ*comR* | NCTC 10919 Δ*comR::cat* (38715…39601) | This study |
| NIDR 6715-7::*comR1* | NIDR 6715-7 pWL52 (*comR1-comS*) | This study |
| NIDR 6715-7::*comR2* | NIDR 6715-7 pWL53 (*comR2*) | This study |
| NIDR 6715-7Δ*comR1-comS* | NIDR 6715-7 Δ*comR1-comS::cat* (38803…39926) | This study |
| NIDR 6715-7Δ*comR2* | NIDR 6715-7 Δ*comR2::cat* (42284…43164) | This study |
| NIDR 6715-7Δ*comR1-comR2* | NIDR 6715-7 Δ*comR1-comR2::cat* | This study |
| NIDR 6715-7::*comR1*Δ*comR1-comS* | NIDR 6715-7 Δ*comR1-comS::cat* pComR1 | This study |
| NIDR 6715-7::*comR2*Δ*comR1-comS* | NIDR 6715-7 Δ*comR1-comS::cat* pComR2 | This study |
| NIDR 6715-7::*comR1*Δ*comR2* | NIDR 6715-7 Δ*comR2::cat* pComR1 | This study |
| NIDR 6715-7::*comR2*Δ*comR2* | NIDR 6715-7 Δ*comR2::cat* pComR2 | This study |
| NIDR 6715-7::*comR1*Δ*comR1-comR2* | NIDR 6715-7 Δ*comR1-comR2::cat* pComR1 | This study |
| NIDR 6715-7::*comR2*Δ*comR1-comR2* | NIDR 6715-7 Δ*comR1-comR2::cat* pComR2 | This study |
| **Plasmids** | | |
| pLacZ | $Spc^r$; pRW17::P23-*lacZ* | This study |
| pComR | $Spc^r$; pRW17::*comR* (from NCTC 10919, 38498…39787) | This study |
| pComR1 | $Spc^r$; pRW17::*comR1-comS* (from NIDR 6715-7, 38585…40006) | This study |
| pComR2 | $Spc^r$; pRW17::*comR2* (from NIDR 6715-7, 42099…43350) | This study |

**Supplementary Table S4. List of Primers**

| **Primer** | **Sequence** | **Note** |
|---|---|---|
| O1-KRMIT-aph3-F | gaaggtctctatagggcaaagcataaaaacttgcatggact | *aph3* cloning |
| O2-KRMIT-aph3-R | gaaggtctctgattacatcagagtatggacagttgcgga | *aph3* cloning |
| O1-cat-F2 | gaaggtctctatagagcttgattttcgttcgtgaatacatgt | *cat* cloning |
| O2-cat-R2 | gaaggtctctgatttcaataatcgcatccgattgcagt | *cat* cloning |
| ***comR* deletion** | | |
| 10-comR-U-F | cttgactcttgagtttgaaaaggctcgt | NCTC 10919 *comR*, NIDR 6715-7 *comR12* deletion upstream arm |
| O1-10-comR-U-R | gaaggtctctctatactgggccctccaatttagtcct | NCTC 10919 *comR*, NIDR 6715-7 *comR12* deletion upstream arm |
| O2-10-comR-D-F | gaaggtctct*aatc*aacgatgggattgagtaagagcgagtt | NCTC 10919 *comR*, NIDR 6715-7 *comR12* deletion downstream arm |
| 10-comR-D-R | tggcacttgaacaaacccttcgt | NCTC 10919 *comR*, NIDR 6715-7 *comR12* deletion downstream arm |
| 67-comR2-U-F | cgtggtcaactctcaatagggcagtt | NIDR 6715-7 *comR2* deletion upstream arm |
| O1-67-comR2-U-R | gaaggtctctctatcgtcataatcgtcctccaattcttggtt | NIDR 6715-7 *comR2* deletion upstream arm |
| comR-check-F2 | aggtggttccctctgggatttca | NCTC 10919 *comR*, NIDR 6715-7 *comR12* deletion check |
| 10comR-check-R2 | tggtcaaccaactagcgacttttca | NCTC 10919 *comR*, NIDR 6715-7 *comR12* deletion check |
| 67-comR2-check-F2 | acccaaccaaaaactcaggtggtca | NIDR 6715-7 *comR2* deletion check |
| cat-i-F | attgtcagataggcctaatgactggct | *comR* deletion check |
| cat-i-R | acctaactctccgtcgctattgtaacca | *comR* deletion check |
| 10comR-o-F | cagctcctactcaagaaggacagaatga | NCTC 10919 *comR*, NIDR 6715-7 *comR12* deletion check |
| 10comR-o-R | tgctacgctttgtagcgcaattcttt | NCTC 10919 *comR*, NIDR 6715-7 *comR12* deletion check |
| 67R2-o-F | agcggcgttactcttcaattgaagg | NIDR 6715-7 *comR2* deletion check |
| ***comR* plasmid complementation** | | |
| pRW17-seq-R | tgagccagtgtgactctagtagaga | Insertion check primer for pRW17 |
| pRW17-seq-F | taccaatgcttaatcagtgagggac | Insertion check primer for pRW17 |
| O1-10comR-F | gaaggtctctatagggtggtatccacgtattggaattcca | NCTC 10919 *comR* cloning |
| O1-67-comR2-F | gaaggtctctatagtccttatagataagacaagcactctcaacaac | NIDR 6715-7 *comR2* cloning |
| O4-10comR-R | gaaggtctctagtccaatggtatcaagagtgagctgacca | NCTC 10919 *comR* cloning |
| ***lox* deletion** | | |
| lox-5-F | acccaatccgtcgagaaattgct | *lox* deletion upstream arm |
| O1-lox-5-R | gaaggtctctctatggactgtcaattatcctgctcagct | *lox* deletion upstream arm |
| O2-lox-3-F | gaaggtctctaatcgtgcccaaaccgttgaagaagt | *lox* deletion downstream arm |
| lox-3-R | agcaatcaaggcaactaggatagca | *lox* deletion downstream arm |
| lox-check-R | agagcctgatggctcccttatca | *lox* deletion check |
| lox-check-F | atttttcttatggagtcgggccagt | *lox* deletion check |

### Supplementary Methods

#### Transformation Protocol for *Streptococcus sobrinus*

1. Inoculate a culture of *S. sobirnus* from a frozen glycerol stock or an agar plate into a 15 ml tube with 10 ml CDM. The CDM should be fresh, as we found that transformation efficiency drops noticeably after CDM has been kept in the fridge for over 3 weeks. Incubate overnight without shaking in an anaerobic chamber at 37°C.

2. At the same time, incubate enough CDM for the next day's dilution in 15 ml or 50 ml tubes with a loosened cap. This serves to prewarm and deoxygenate the media.

3. Dilute the overnight culture 150-fold into a culture tube with the pre-incubated CDM. The higher dilution seems to increase transformation efficiency. Incubate anaerobically at 37°C while monitoring the $OD_{600}$ with a portable cell density meter.

4. When the $OD_{600}$ reaches about 0.6, add XIP (LMCTIAR for NIDR 6715-7 or LMCTIVR for SL1 dissolved in DMSO at 10 mM) to a final concentration of 10 µM. Mix well by vigorously shaking the culture tube.

5. Immediately aliquot 200 µl of the culture into a microcentrifuge tube. Add appropriate amount of DNA (as a reference, 400 ng purified PCR product was used for our strain characterization experiments, but efficiency varies depending on the DNA sequence). Mix well by flicking the tube 15 times. (We found that flicking produces noticeably better results than mixing with a pipette; for this reason, we use 500 µl tubes instead of 1.5 ml tubes as the former are easier to flick.)

6. Incubate the capped tubes for 2 h.

7. Prewarm two THY agar plates with appropriate antibiotic selection.

8. Plate 150 ul of the culture onto a selection plate. Plate a small amount of culture that did not have added DNA to another selection plate as a negative control.

9. Incubate the plates at 37°C in a 5% $CO_2$ chamber for 24 h or until colonies are visible.

### Plasmid Construction

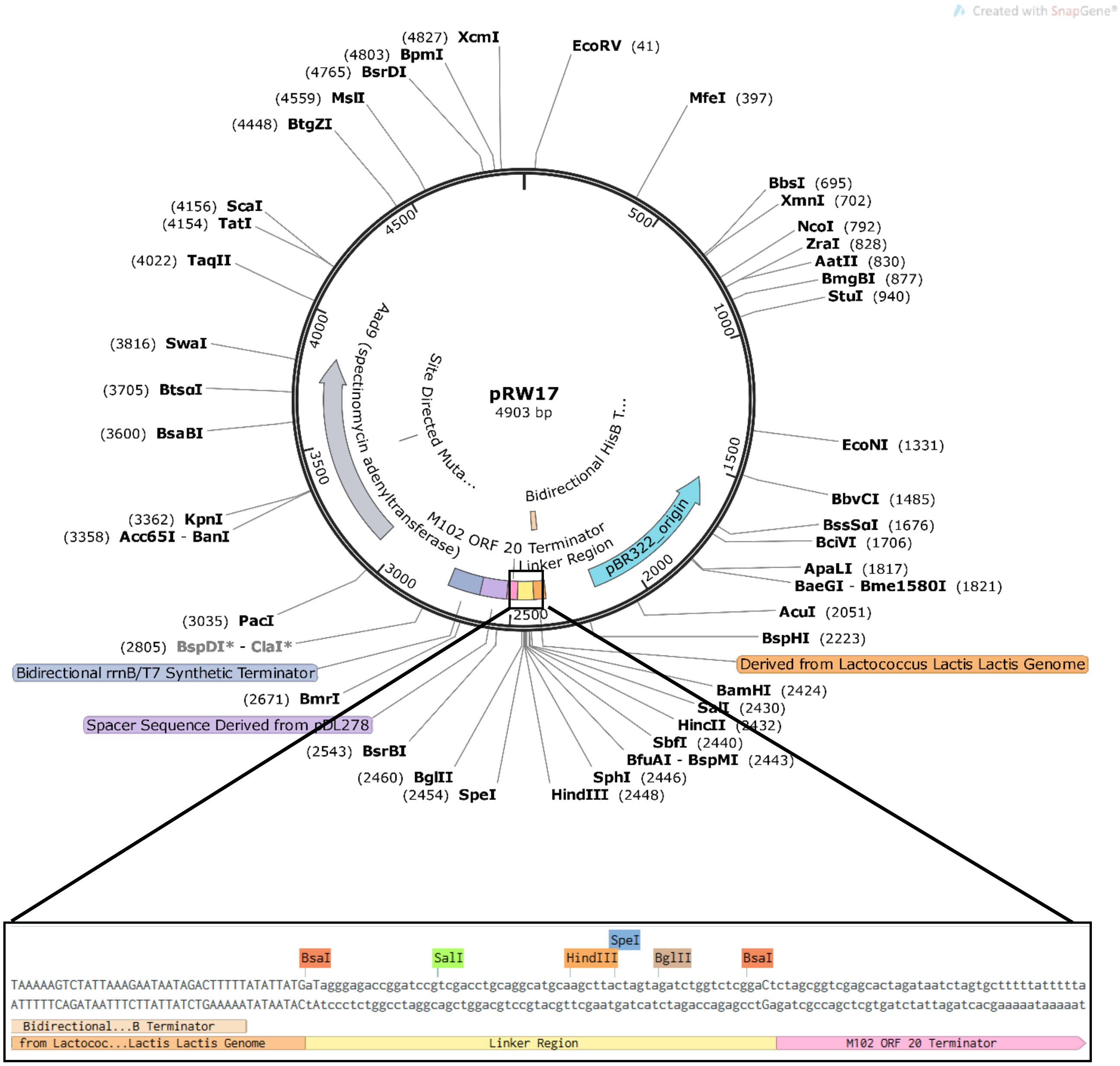


**Supplementary Figure S1.** pRW17 plasmid map and linker region.

pRW17 is a streptococcal expression vector designed to enable Golden Gate assembly of modular gene expression cassettes. The backbone of pRW17 was derived from pRW06, a derivative of pZX10 (Xie et al. 2013) with the Gram-negative origin of replication from pDL278 (LeBlanc et al. 1992) inserted into the BbvCI site. pRW17BB.for (5'-cggatgggtctcggagtagaagatcgattttcgttcg-3') and pRW17BB.rev (5'-tgctcgggtctccgtccctcactgattaagcattgg-3') were used to perform PCR (Q5 Hot Start High Fidelity, NEB) using pRW06 as template. The resulting amplicon contained terminal 5' and 3' BsaI restriction sites flanking the *aad9* spectinomycin resistance marker, the Gram-positive origin of replication, and the Gram-negative origin of replication. The insert molecule for the construction of pRW17 was designed to include the several components and synthesized (GeneBlock, IDT). A polylinker including restriction sites for many commonly utilized restriction enzymes, including BsaI, was included between two intrinsic, transcriptional terminators. The *hisB* bidirectional terminator from the histidine biosynthesis cluster of *Lactococcus lactis* subsp. lactis (Delorme et al. 1992) was placed on the 5' flank of the polylinker, while a predicted, unidirectional terminator associated with ORF 20 of the *Streptococcus mutans* UA159 bacteriophage M102 (van der Ploeg , 2007) was placed in series with the synthetic bidirectional rrnB T1/T7-TE terminator on the 3' flank (Glass and Riedel-Kruse 2018). The synthesized insert was amplified upon arrival with PCR using the following primers: Insert.GG.Comp.for (5'-gtatacgaagacacggactacaacagaccttgct -3') and Insert.GG.Comp.rev (5'-acggctgaagacacactcctttgtttatcctcctc -3'). The resulting amplicon had BbsI sites on the 5' and 3' end, which upon digestion yield overhangs compatible with the overhangs on the backbone produced from BsaI digestion. Each component was digested with its respective enzyme separately and then utilized in a standard T4 DNA Ligase-mediated ligation reaction using 40 fmol of digested insert and 20 fmol of digested backbone. After 1 hour, 5 uL of the 20 uL ligation mix was used to transform competent E. coli DH5a (Mix & Go, Zymo Research). After a 1-hour outgrowth, the cells were plated on LB/Spec100 and incubated overnight at 37 C. The next morning, colonies were screened with colony PCR. Subsequent plasmid isolation, restriction enzyme digest and Sanger sequencing validated successful pRW17 construction.

For use as a Golden Gate cloning platform a set of five, orthogonal four-nucleotide overhangs were developed by surveying previous literature using similar designs (Whitaker et al. 2017) and by referencing the T4 ligation fidelity and efficiency overhang profile developed by Potapov et al. 2018.

```
Overhang 1:  5'-*ATAG -3'
             3'- TATC*-5'

Overhang 2:  5'-*TCTA -3'
             3'- AGAT*-5'

Overhang 3:  5'-*AATC -3'
             3'- TTAG*-5'

Overhang 4:  5'-*GACT -3'
             3'- CTGA*-5'

Overhang 5:  5'-*CATA -3'
             3'- GTAT*-5'
```
